# Time-series trend of pandemic SARS-CoV-2 variants visualized using batch-learning self-organizing map for oligonucleotide compositions

**DOI:** 10.1101/2021.04.15.439956

**Authors:** Takashi Abe, Ryuki Furukawa, Yuki Iwasaki, Toshimichi Ikemura

## Abstract

To confront the global threat of coronavirus disease 2019, a massive number of the severe acute respiratory syndrome coronavirus 2 (SARS-CoV-2) genome sequences have been decoded, with the results promptly released through the GISAID database. Based on variant types, eight clades have already been defined in GISAID, but the diversity can be far greater. Owing to the explosive increase in available sequences, it is important to develop new technologies that can easily grasp the whole picture of the big-sequence data and support efficient knowledge discovery. An ability to efficiently clarify the detailed time-series changes in genome-wide mutation patterns will enable us to promptly identify and characterize dangerous variants that rapidly increase their population frequency. Here, we collectively analyzed over 150,000 SARS-CoV-2 genomes to understand their overall features and time-dependent changes using a batch-learning self-organizing map (BLSOM) for oligonucleotide composition, which is an unsupervised machine learning method. BLSOM can separate clades defined by GISAID with high precision, and each clade is subdivided into clusters, which shows a differential increase/decrease pattern based on geographic region and time. This allowed us to identify prevalent strains in each region and to show the commonality and diversity of the prevalent strains. Comprehensive characterization of the oligonucleotide composition of SARS-CoV-2 and elucidation of time-series trends of the population frequency of variants can clarify the viral adaptation processes after invasion into the human population and the time-dependent trend of prevalent epidemic strains across various regions, such as continents.

## INTRODUCTION

The severe acute respiratory syndrome coronavirus 2 (SARS-CoV-2) has spread rampantly worldwide since it was first reported in December 2019, and its momentum is still ongoing (WHO. 2020). To address the SARS-CoV-2 pandemic in detail, genome sequencing has been performed on a global scale and published by GISAID (Elbe et al. 2017), the SARS-CoV-2 genome database, having more than 780,000 viral sequences as of March 2021 (https://www.gisaid.org/). SARS-CoV-2 is an RNA virus with a fast evolutionary rate that has already been classified into eight clades by GISAID, and epidemics caused by new variant have been known to occur (Benvenuto et al. 2020; Gorbalenya et al. 2020; Sun et al. 2020; Hu et al. 2021; Kirby 2021; Wang et al. 2021). Because the number of registered genome sequences is increasing explosively, it has become difficult to cope with the current and future situation using only the conventional phylogenetic tree method based on multiple sequence alignment, which requires an enormous amount of computation time for a massive number of sequences. Therefore, it is imperative to develop a sequence alignment-free method that will enable us to easily grasp the whole picture of the big-sequence data and support efficient knowledge discovery from it.

By focusing on the frequency of short oligonucleotides (e.g., tetra- and penta-nucleotides) in a large number of genomic fragments (e.g., 10 kb) derived from a wide variety of species, we have developed an unsupervised explainable AI (batch-learning self-organizing map; BLSOM), which enables separation (self-organization) of the genomic sequences by species and phylogeny and explains the causes that contribute to this separation (Abe et al. 2003). In the analysis of genomic fragments of a wide range of microbial genomes, over 5 million sequences can be separated by phylogenetic groups with high accuracy (Abe et al. 2020).

In a prior analysis of all influenza A strains, viral genomes were separated (self-organized) by host animals based only on the similarity of the oligonucleotide composition, although no host information was provided during BLSOM learning (Iwasaki et al. 2011). On a single map, all viral sequences could be separated, and notably, BLSOM is an explainable AI that can explain diagnostic oligonucleotides, which contribute to host-dependent clustering. When studying the 2009 swine-derived flu pandemic (H1N1/2009), we could detect directional time-series changes in oligonucleotide composition because of possible adaptations to the new host, namely humans (Iwasaki et al. 2011), showing that near-future prediction was possible, albeit partially (Iwasaki et al. 2013).

We have previously revealed lineage-specific oligonucleotide compositions for a wide range of virus lineages and established a method to identify and classify viral-derived sequences in tick intestinal metagenomic sequences (Qiu et al. 2019). In the case of SARS-CoV-2, we analyzed time-series changes in mono- and oligo-nucleotide compositions and found their time-dependent directional changes that are thought to be adaptive for growth in humans, which allowed us to predict candidates of advantageous mutations for growth in human cells (Ikemura et al. 2020; Wada, Wada & Ikemura. 2020; Iwasaki Abe & Ikemura. 2021). Furthermore, we recently performed BLSOM analysis on di-to penta-nucleotide compositions in approximately 150,000 SARS-CoV-2 genomes. Because the accuracy of separation by clade increased as the oligonucleotide length increased, in this report, we present the BLSOM results for the pentanucleotide composition. BLSOM could serve as a powerful tool for elucidating comprehensive characterization of the oligonucleotide composition of SARS-CoV-2 and time-series trends of prevalent epidemic strains across various regions, such as continents.

## METHODS

### SARS-CoV-2 genome sequences

The full-length genome sequences of SARS-CoV-2 were downloaded from the GISAID database on November 4, 2020. The total number of sequences was 170,190. From these sequences, those with a length of more than 27 kb after removing the polyA-tail sequences were selected.

### Oligonucleotide frequency and odds ratio

Pentanucleotide frequencies and odds ratios were used in the present study. The pentanucleotide odd ratios (observed/expected values) were calculated using the formula P_VWXYZ_ = *f*_VWXYZ_/*f*_V_*f*_W_*f*_X_*f*_Y_*f*_Z_, where *f*_V_, *f*_W_, *f*_X_, *f*_Y_ and *f*_Z_ denote the frequencies of mononucleotides V, W, X, Y and Z, respectively, and *f*_VWXYZ_ denotes the frequency of pentanucleotide VWXYZ (Karlin et al. 1998).

### BLSOM

Kohonen’s self-organizing map (SOM), an unsupervised neural network algorithm, is a powerful tool for clustering and visualizing high-dimensional complex data on a two-dimensional map (Kohonen, 1990; Kohonen et al., 1996). We modified the conventional SOM for genome informatics on the basis of batch learning, aiming to make the learning process and the resulting map independent of the order of data input (Kanaya et al. 2001; Abe et al. 2003). The newly developed SOM, BLSOM, is suitable for high-performance parallel computing and, therefore, for big data analysis. The initial weight vectors were defined using principal component analysis (PCA), based on the variance-covariance matrix, rather than by using random values. The weight vectors (wij) were arranged in a two-dimensional lattice denoted by i (= 0, 1,…, I-1) and j (= 0, 1,…, J-1) and were set and updated as described previously (Kanaya et al. 2001; Abe et al. 2003). A BLSOM program suitable for PC cluster systems is available on our website (http://bioinfo.ie.niigata-u.ac.jp/?BLSOM).

## RESULTS and DISCUSSION

### BLSOMfor pentanucleotide composition and their odds ratio

It should be mentioned here that SARS-CoV-2 genomes have changed their mononucleotide composition during the course of the epidemic in humans, reducing C and increasing U, regardless of clade (Mercatelli et al. 2020; Wada, Wada & Ikemura. 2020; Iwasaki Abe & Ikemura 2021), a process which is thought to be caused by the APOBEC family enzymes (Mangeat et al. 2003; Simmonds 2020). Considering this clade-independent tendency, we performed BLSOM analysis of not only the pentanucleotide composition but also their odds ratio, which can reduce the effects caused by changes in the mononucleotide composition. Additionally, to check the robustness of sequence accuracy, we used datasets with different sequence accuracies: 167,905 sequences with less than 10% unknown nucleotides other than ATGCs in the genome sequence and 130,753 sequences with less than 1% unknown nucleotides; for each sequence dataset, the number of cases by region and clade is shown in Table 1.

**Table 1.**
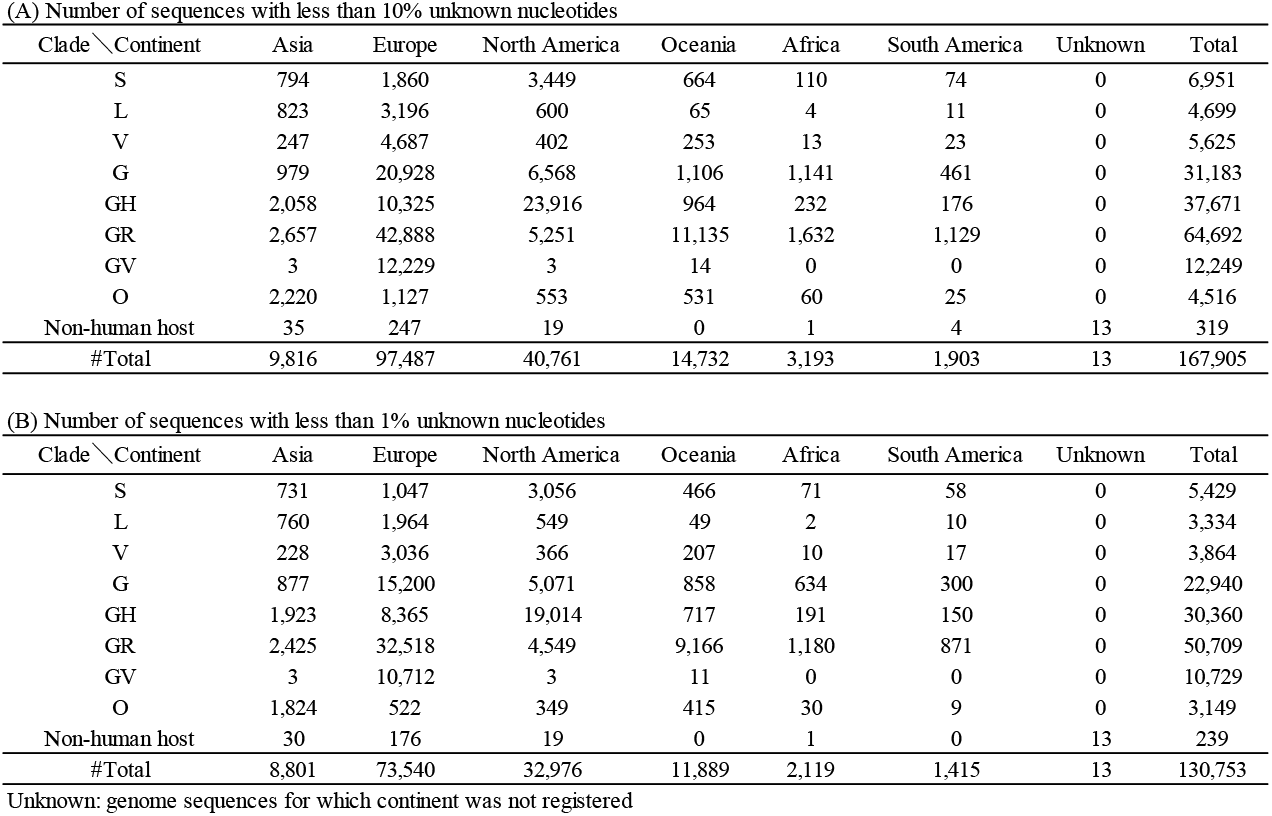
Number of SARS-CoV-2 genome sequences with less than 10% (A) and less than 1% (B) unknown nucleotides used in this study.

First, we constructed BLSOM for sequences with less than 10% unknown nucleotides, using the pentanucleotide composition and their odds ratios (Figure 1A and B). BLSOM utilizes unsupervised machine learning, and the genome sequences are clustered (self-organized) on a two-dimensional plane, based only on the difference in the vector data in a 1024 (=4^5^)-dimensional space. Lattice points that include sequences from more than one clade are indicated in black, those that contain no genomic sequences are indicated by blank, and those containing sequences from a single clade are indicated in the color representing the clade. The odds ratio (Figure 1B) gave more accurate separations (a smaller percentage of black grid points), possibly by excluding effects owing to the clade-independent time-series change in the mononucleotide composition (Iwasaki Abe & Ikemura. 2021), which affected all SARS-CoV-2 clades. Even for the sequences with low-sequence accuracy, clade-dependent separation occurs, allowing us to understand characteristics of the oligonucleotide composition that are specific to each clade; thus, oligonucleotide-BLSOM is thought to be a robust method. However, it is clear that BLSOMs for sequences with less than 1% unknown nucleotides (Figure 1C and D) gave more accurate separation than those listed in Figure 1A and B, and the highest resolution was obtained for the BLSOM for the odds ratio (Figure 1D).

**Figure 1.**
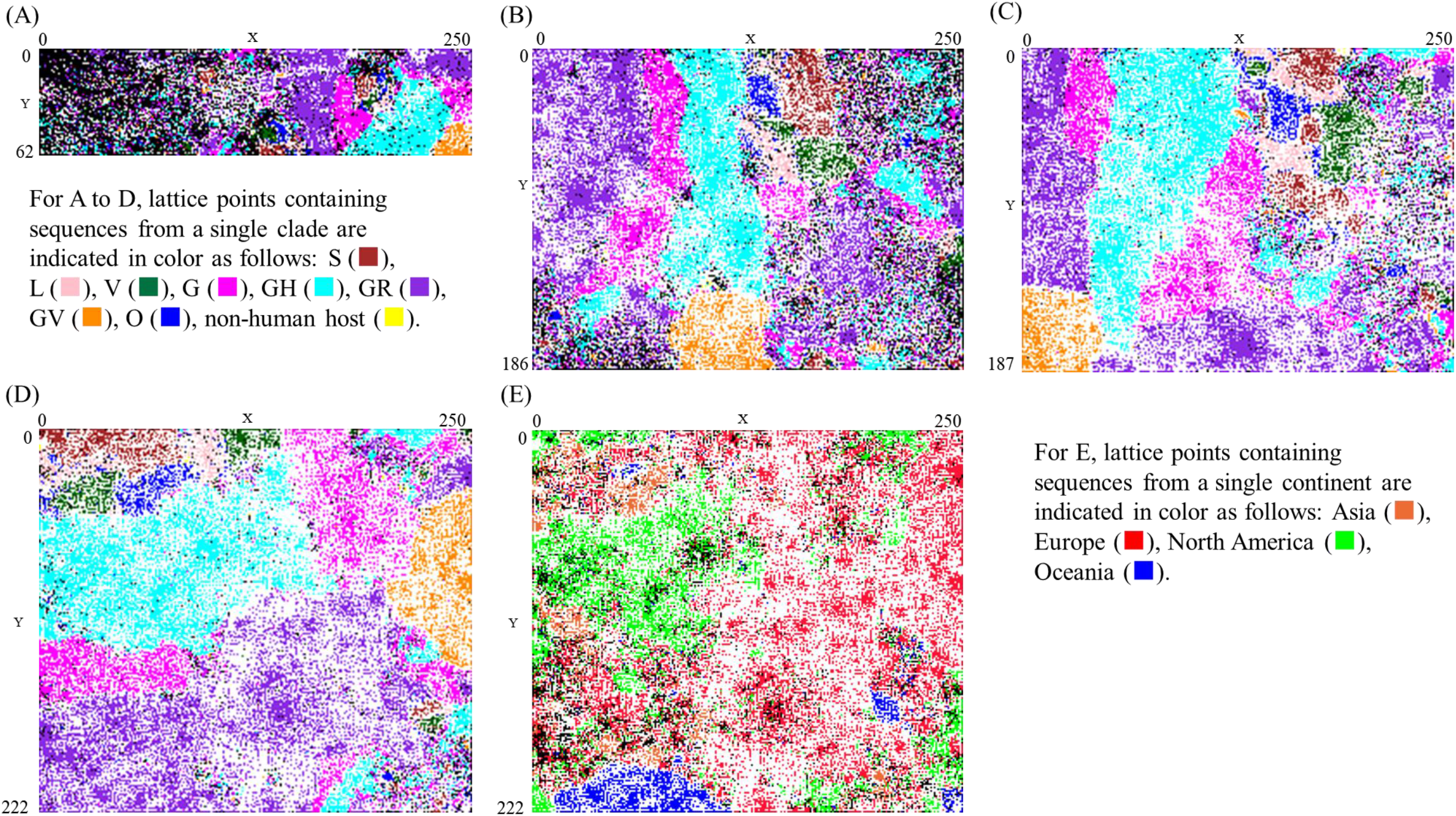
BLSOM for pentanucleotide usage. (A) Pentanucleotide composition and (B) their odds ratio for sequences with less than 10% unknown nucleotides. (C) Pentanucleotide composition and (D) their odds ratio for sequences with less than 1% unknown nucleotides. Lattice points that include sequences from more than one clade are indicated in black, those that contain no genomic sequences are indicated by blank, and those containing sequences from a single clade are indicated in color as follows: S 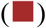, L 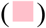, V 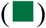, G 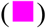, GH 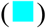, GR 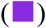, GV 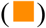, O 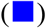, non-human host 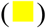. (E) Distribution of sequences by continent on the BLSOM with the pentanucleotide odds ratio. Lattice points that include sequences from more than one continent are indicated in black, those that contain no genomic sequences are indicated by blank, and those containing sequences from a single continent are indicated in color as follows: Asia 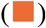, Europe 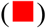, North America 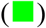, Oceania 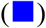.

Clades have been defined by the statistical distribution of phylogenetic distances in tree construction based on multiple sequence alignments (Han et al. 2019; Tang et al. 2020), whereas BLSOM is a sequence alignment-free analysis that is suitable for the analysis of massive data. Because sequences at different locations on BLSOM have different oligonucleotide compositions, clustering according to clades means that sequences belonging to different clades have different oligonucleotide combinations, that is, differential combinations of mutations.

### 3D display of the data for different continents

Using BLSOM (Figure 1D) for the pentanucleotide odds ratio, Figure 1E examines the classification according to four continents (Asia, Europe, North America, and Oceania) that have large numbers of sequences. Here, the lattice points containing sequences of different continents are displayed in black, and those containing only sequences of a single continent are displayed in the color specifying each continent. Although not as clear as clade-dependent separations, regional differences have been observed, which should reflect differential shares of prevalent variants among continents. However, it is apparently difficult to obtain sufficient information from the results shown in Figure 1E alone. BLSOM is equipped with various visualization tools for analysis results; therefore, we next show the number of sequences belonging to each lattice point with a 3D display.

Again, using the BLSOM shown in Figure 1D, Figure 2 shows the number of sequences belonging to each lattice point for each clade in each continent as a vertical bar, which is colored by continent, as shown in Figure 1E. Looking laterally at a particular clade, each clade consists of several subclusters, each consisting of several high peaks surrounded by many low peaks. Different subclusters observed in each clade are distinguished by numbering in each figure, but if they are located in the same zone on BLSOM, the same number is given even if they are of different continents. Looking vertically at a particular continent, sequences of different subclusters of different clades exist in different amounts, and some subclusters are only in a particular continent, that is, the prevalent variants for each continent can be visualized in an easy-to-understand manner. In Supplementary Figure S1, the data shown in Figure 2 are displayed in 2D, and referring to the quantitative results in Figure 2, we defined sequences attributed to each subcluster in each clade.

**Figure 2.**
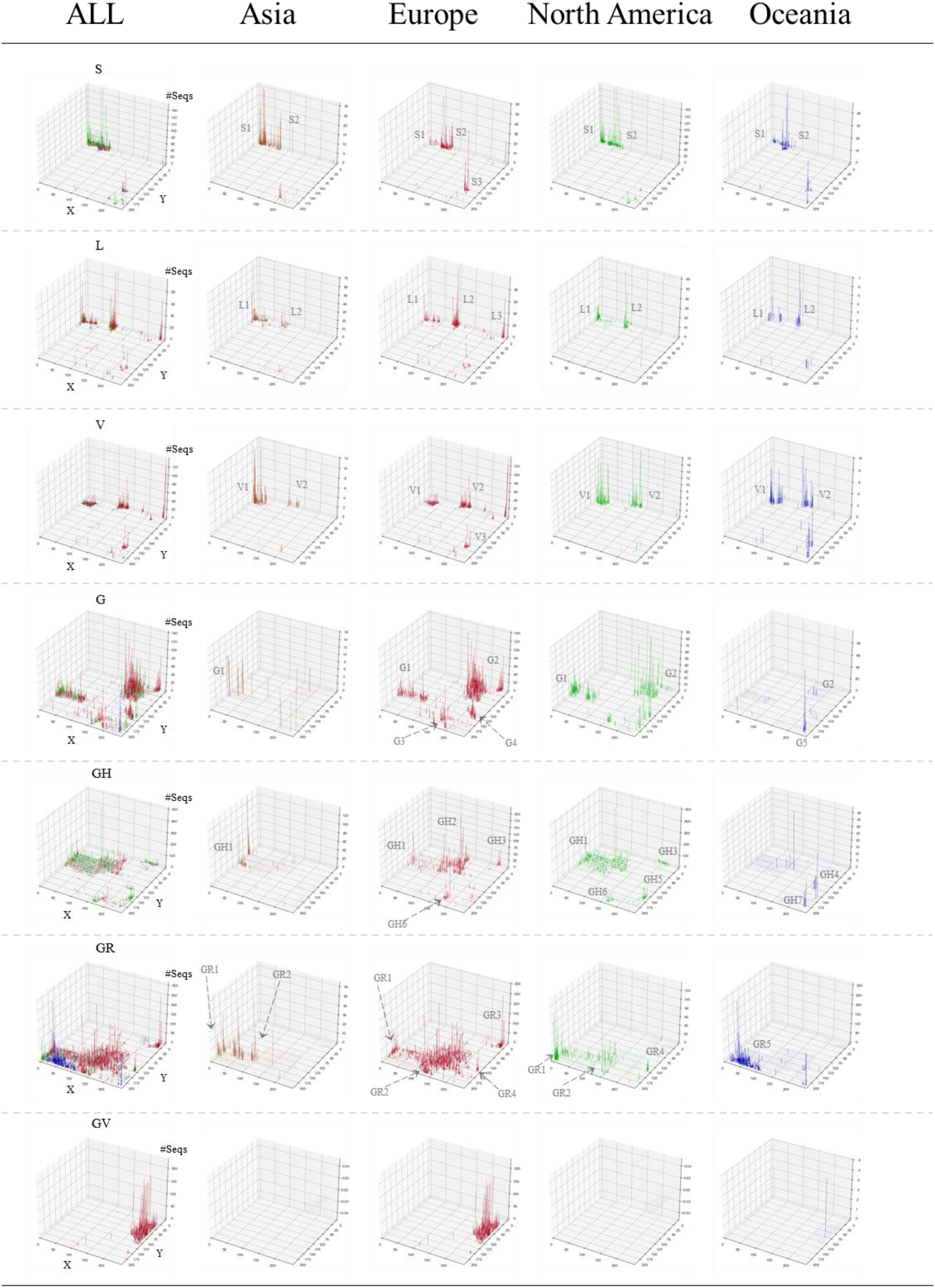
3D display of viral classification by clade and continent. The Z-axis corresponds to the number of sequences attributed to each lattice point. Results for all continents are shown in the ALL panel for each clade. In clades G, GH, GR and GV, lattice points where less than 5 sequences exist are not shown. The vertical bars for individual continents are distinguished by the following colors: Asia 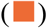, Europe 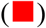, North America 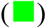, Oceania 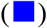. Different subclusters are given suffix numbers.

### Time-series analysis

The fact that sequences belonging to one clade were clearly separated on BLSOM indicates the importance of subdivision of each clade, and the separation on BLSOM is thought to be a good indicator of this subdivision. To further examine the biological significance of the subclusters of each clade on BLSOM, we visualized the number of sequences collected in each month in each region as a vertical bar differentially colored according to clade (Figure 3). Looking laterally at a continent, the time-series quantitative changes among different clades or different subclusters of one clade are clear. Looking at the results for a particular collection month for different continents longitudinally, quantitative changes among different clades or different subclusters of one clade are again clear, depending on the continent.

**Figure 3.**
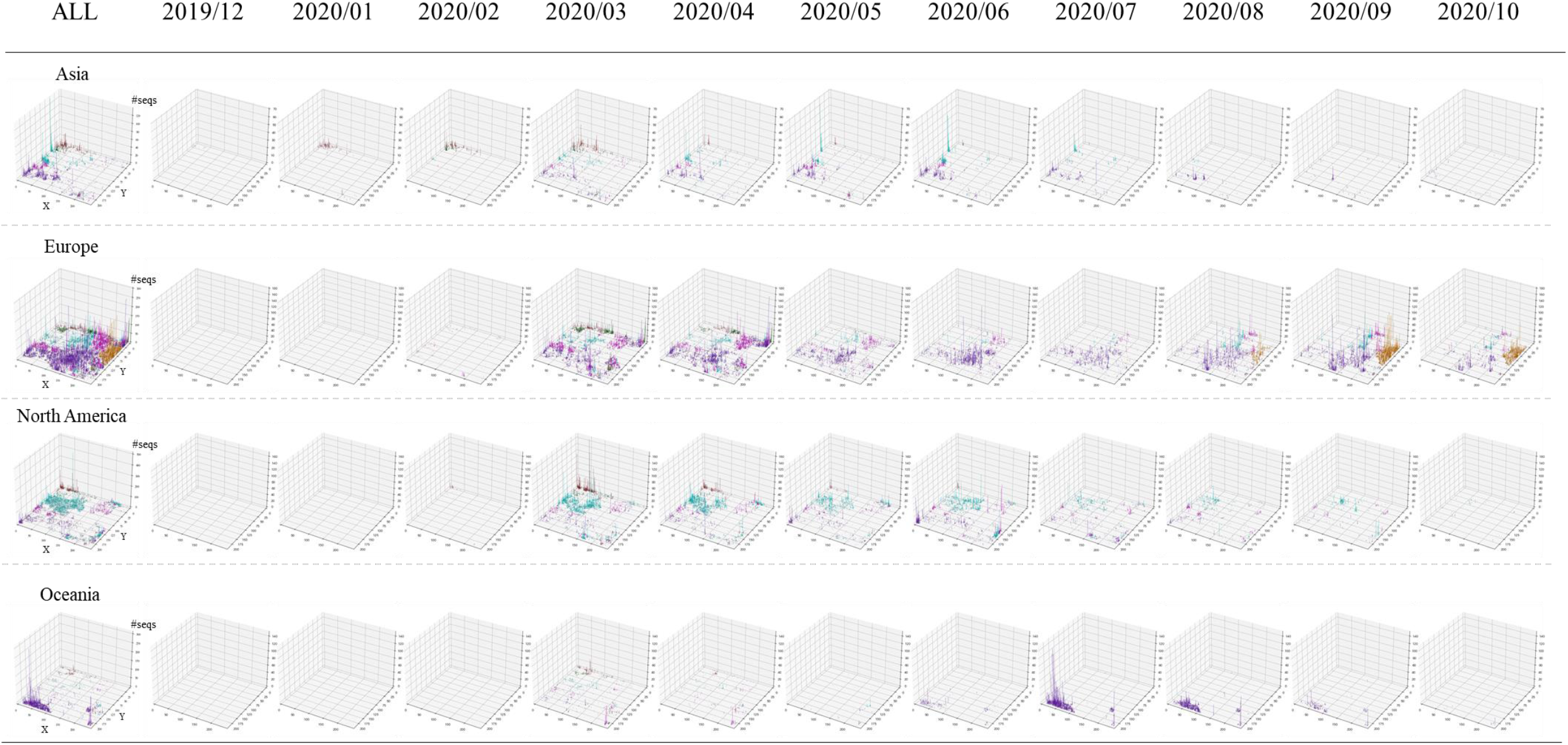
3D display of temporospatial changes. The Z-axis corresponds to the number of sequences attributed to each lattice point. Results for all collection months are shown in the ALL panel for each continent. The vertical bars for individual clades are distinguished by the following colors: S 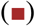, L 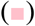, V 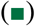, G 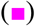, GH 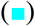, GR 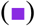, GV 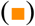.

Next, for each clade in each continent, we quantitatively analyzed the time-series changes in the proportion of its subclusters using a 100% stack bar graph (Figure 4). The percentage of sequences in different subclusters are distinguished by different colors, and when a total number of sequences for a certain month is more than 100, the data for that month is indicated by a thick horizontal bar. We focused mainly on such months.

**Figure 4.**
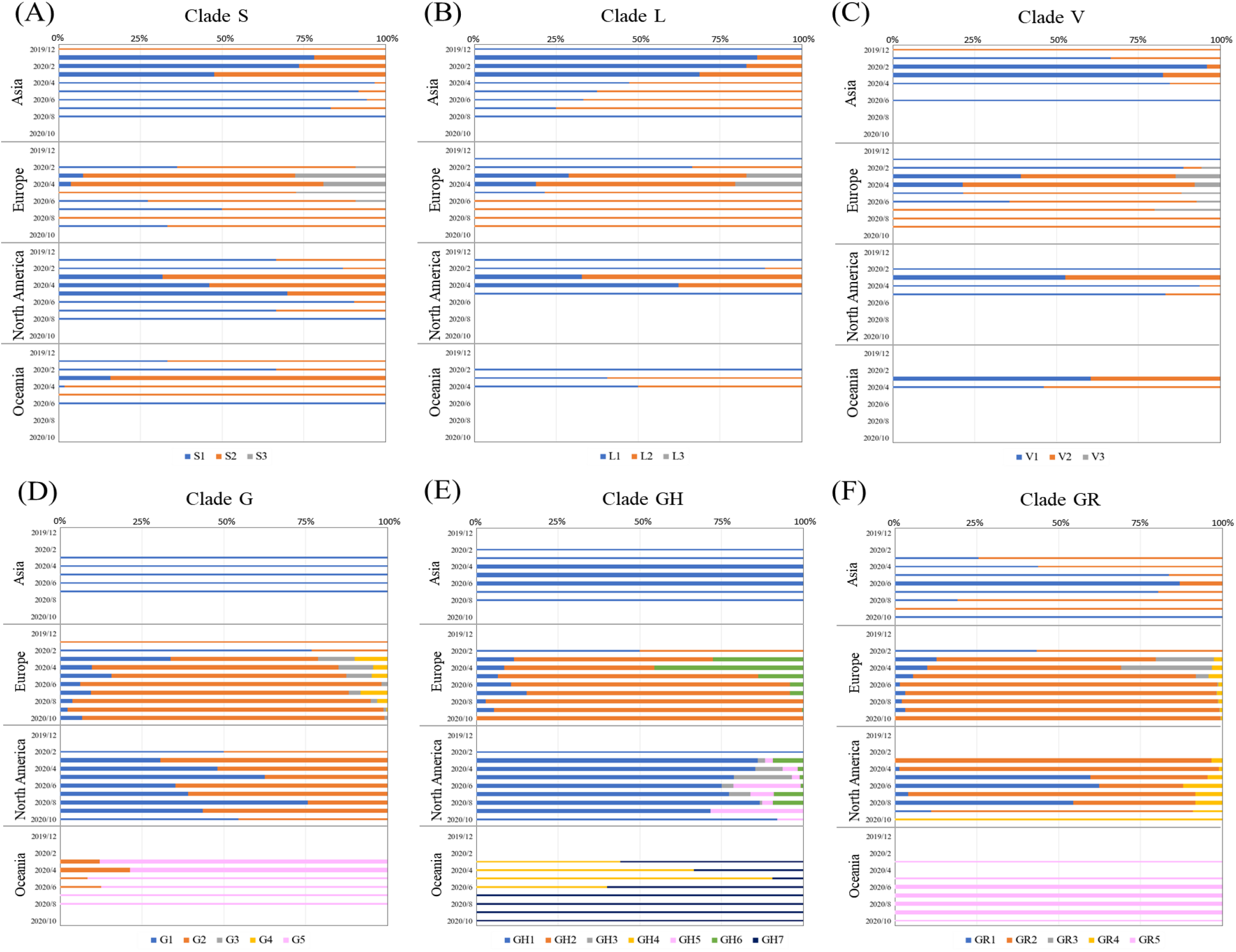
Analysis of 100% stack bar graph for time-series transition in each continent for each subcluster in clades S (A), L (B), V (C), G (D), GH (E) and GR (F). The colors of each subcluster are indicated at the bottom of each figure. The results for months with more than 100 sequences are shown as thick horizontal bars. The number of sequences used in this analysis is given in Supplementary Table S1.

In the clade S/L/V detected in the early stage of the epidemic (December 2019-March 2020), three major subclusters of each clade were observed and distinguished by suffix numbers, and most sequences belonged to the two subclusters: S1/L1/V1 and S2/L2/V2. In Asia, many sequences belonging to S1/L1/V1 were detected in December 2019, but in Europe and other regions, S2/L2/V2 were more abundantly detected in March and April 2020 than S1/L1/V1, and the proportion became more pronounced in April than in March. In March and April in Europe, a remarkable number of sequences belonging to S3/L3/V3 were also detected, showing three different variants prevalent at the beginning of the epidemic in Europe. Far fewer than 100 sequences were detected after May; sequences belonging to S1/L1/V1 were mainly detected in Asia and those belonging to S2/L2/V2 were shown in other regions, presenting differential trends in prevalent variants among continents.

For clade G, which started the epidemic in Europe in February, we defined five subclusters. In February, roughly equal amounts of sequences belonging to G1 and G2 were detected in Europe and North America, but as the epidemic progressed, those belonging to G2 were mainly detected in Europe, whereas those belonging to both G1 and G2 were prevalent in North America. In Asia, only sequences belonging to G1 are detected; in Oceania, those belonging to G2 accounted for about 10% in the early stage, but afterward, those belonging to Oceania-specific G5 accounted for the majority.

For GH, we defined seven subclusters, including GH1 and GH2, which dominated in North America and Europe, respectively. In North America, in addition to GH1, several months contain approximately 20% of the sequences belonging to GH3, GH5, and GH6. In Asia, only GH1 has been detected. In Oceania, only GH4 and GH7, which were specific to this region, were detected; initially, GH4 was dominant, but after July, GH7 was primarily detected.

For GR, we defined five subclusters, including GR1 and GR2, which dominated in North America and Europe, respectively. Moreover, in Europe, GR1 was detected to the same extent as GR2 in February, but as the epidemic progressed, GR2 began to predominate. In North America, the occupancy of GR1 and GR2 varied to some extent depending on the collection month. In Asia, GR1 was mainly detected, and in Oceania, only region-specific subclusters have been detected.

These temporospatial changes in subclusters show that the subcluster is the separation (self-organization) that reflects biological significance and is fundamental information for understanding the overall picture of the SARS-CoV-2 variants.

## CONCLUSION and PERSPECTICES

Based on the phylogenetic tree construction by multiple sequence alignments, GISAID has defined seven clades of SARS-CoV-2, giving a total of eight if clade O corresponding to others is included. However, these classifications are clearly inadequate to understand the current status of SARS-CoV-2 because this RNA virus evolves at a high speed. Using only the oligonucleotide composition of many genomic sequences, the unsupervised machine learning, BLSOM, could separate viral sequences according to not only clades but also subclusters within each clade. The separation (self-organization) that AI can accomplish without any hypothesis or model is thought to be a classification from a new perspective. BLSOM is equipped with various tools that allow us to visualize the analysis results in an easily understandable way and to visualize differences in the number of subcluster sequences among continents (Figure 2) and their time-series changes (Figure 3), i.e., the distinct variations in the resulting subclusters depending on the region and the collection time.

Herein, we focused on pentanucleotide composition, but similar separations were obtained for other lengths of oligonucleotides (Ikemura et al. 2020). BLSOM is an explanatory AI that can clarify combinatorial patterns of oligonucleotides that contribute to the separation according to clades and their subclusters. BLSOM is a powerful method for elucidating comprehensive characterization of the oligonucleotide composition in a massive number of SARS-CoV-2 genome sequences. Next, it will be important to know the relationship between the strains isolated in clades and their subclusters and the causative mutations. When it comes to oligonucleotides as long as 15-mers, most are only present in one copy in the viral genome; therefore, changes in 15-mer sequences can be directly linked to mutations, and we have already started analysis from this perspective (Ikemura et al. 2020). The implementation of time-series oligonucleotide analysis of variants with rapidly expanding intrapopulation frequencies has enabled the identification of candidates for advantageous mutations for viral infection and growth in human cells (Wada, Wada & Ikemura. 2020).

Phylogenetic methods based on sequence alignment have been widely used in evolutionary studies (Hadfield et al. 2018; Kumar et al. 2018), and these methods are undoubtedly essential for studying the phylogenetic relationships between different viral species and variations in the same virus at the single-nucleotide level. In contrast, AI can analyze a massive number of SARS-CoV-2 sequences at once without difficulty, potentially reaching a level of one million in the near future. The AI method for oligonucleotide composition has become increasingly important as a complement to the phylogenetic tree construction method in preparing for future outbreaks of various infectious RNA viruses.

## AUTHORS’ CONTRIBUTIONS

TA and TI conceived and designed research; TA, RF, and TI performed research; TA, RF, and YI analyzed data; and all authors wrote the paper.

## ACKNOWLEDGEMENTS

We gratefully acknowledge the authors submitting their sequences from GISAID’s Database.

## SUPPLEMENTARY DATA

**Supplementary Figure S1.**
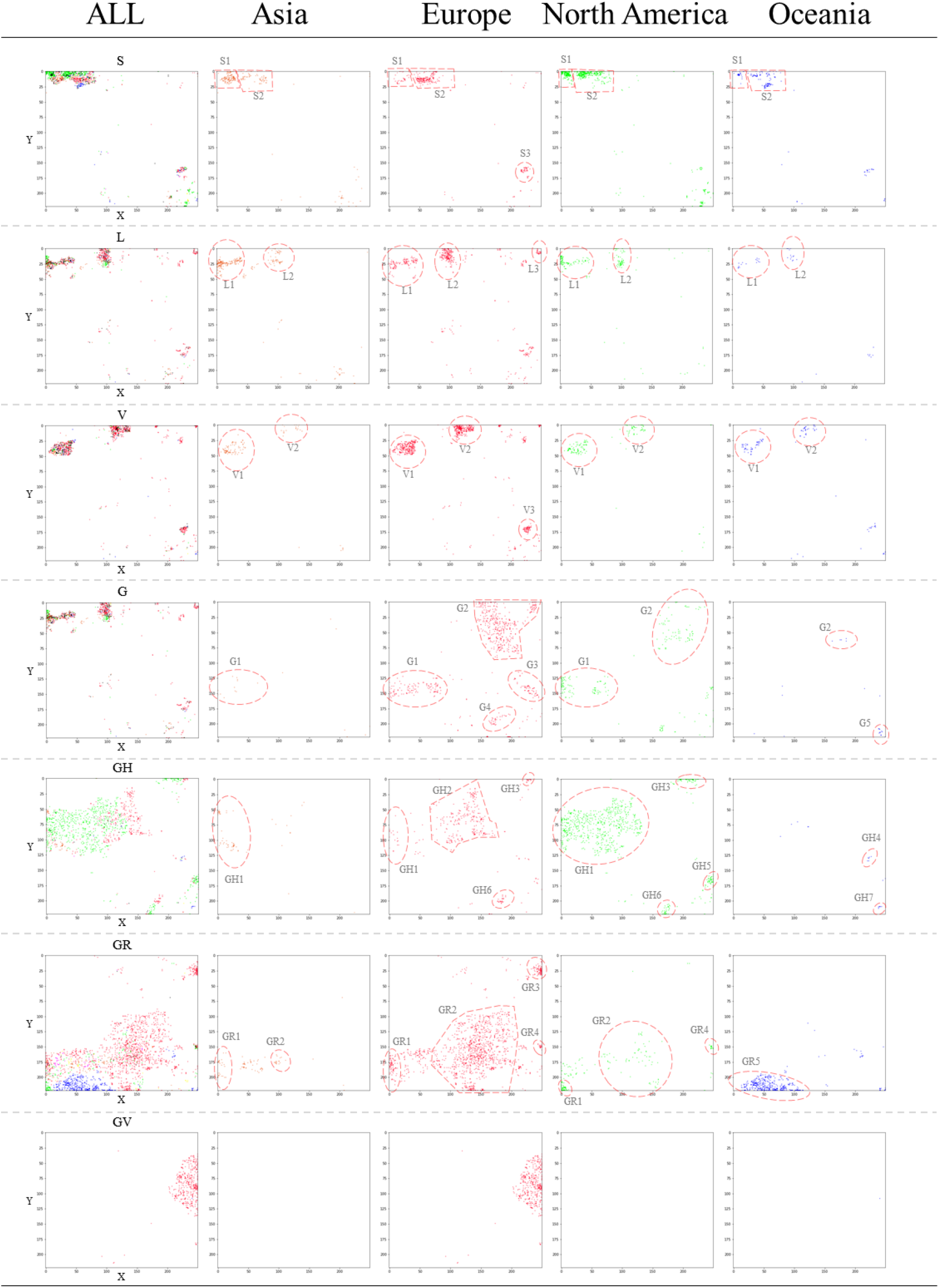
2D display of the classification by clade and continent shown in Figure 2. Each subcluster territory is circled by a dotted line. In clades G, GH, GR and GV, lattice points where less than 5 sequences exist are not shown. The sequences belonging to each territory defined here are used for the analysis in Figure 4.

**Supplementary Table S1.**
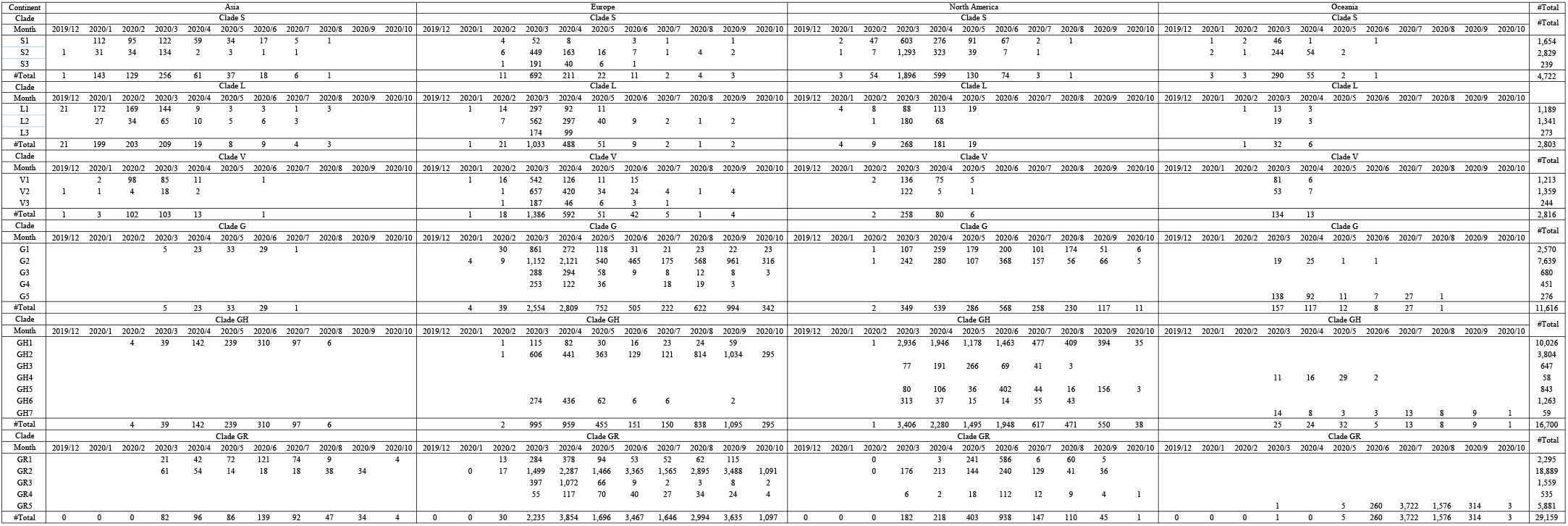
Sequence number of subdivided clusters in clade for each month by continent.

